# Co-stimulatory molecules decide T cell fate through regulations of their invigoration and impairment

**DOI:** 10.1101/2022.09.21.508809

**Authors:** Kenji Ichikawa, Kazuhisa Ichikawa

## Abstract

T cell invigoration is an essential step for eliminating pathogens and cancer cells. Co-stimulatory molecules, such as CD80, CD86, and ICOSLG reinforce TCR stimuli for T cell activations. Despite identifying multiple co-stimulatory molecules, the differences of those in downstream signaling have remained unclear. Here, we unravel the differences in avidity of co-stimulatory molecules with T cells cause distinct T cell fates. Specially, CD80 + TCR stimulus promotes induction of multiple T cell effector genes based on prolonged and magnified activation of ERK and AKT compared with other combined stimuli. Long-term and robust activation of these signaling pathways leads to T cell impairment by induction of PD1, and exhausted T cells are vulnerable to disrupt effector functions by interactions with PDL1. Collectively, we reveal the quantitative differences in binding activities of co-stimulatory molecules to T cells cause qualitative differences in downstream signals and gene expressions, thereby branching T cell fates.

## Introduction

Co-stimulations are indispensable for adequate invigorations of T cells (Jenkins et al., 1991; Mueller et al., 1989). In case of the loss of co-stimulations, T cell activations are insufficient, resulting in dysfunctions for eliminating pathogens and tumor cells and causing T cell anergy (Fathman and Lineberry, 2007; Harding et al., 1992). CD28 and ICOS are the representative co-stimulatory molecules on T cells and are closely related genes, but the functions of these two receptors are non-redundant (Esensten et al., 2016). Two CD28 ligands, CD80 and CD86 are co-stimulatory molecules and show restricted expression profiles (Khayyamian et al., 2002; Linsley et al., 1991a). By contrast, ICOSLG shows wide expression profile (Nurieva et al., 2003). It is assumed that the functional differences are yielded by expression profiles of co-stimulatory molecules that bind to CD28 or ICOS (Esensten et al., 2016; Linterman et al., 2009). However, difference in activated pathway and gene expressions under the co-stimulations remained unclear for a long time.

PDL1 and PDL2 disrupts the T cell activations through engagement with PD1 (Okazaki and Honjo, 2007). PD1 negatively regulates TCR and CD28 signaling by dephosphorylations through SHP2 of key molecules in the pathway, thereby suppressing effector activities (Hui et al., 2017; Yokosuka et al., 2012). PD1 preferentially suppress gene expressions of cytokines and effector molecules rather than transcriptional factors (Shimizu et al., 2020). Although PD1 also inhibit gene expressions in lower-affinity T cells with peptide-MHC complexes (Shimizu et al., 2021), the involvement of co-stimulatory molecules in the suppressions of T cell activations is not completely clear.

Despite the critical role of the co-stimulatory molecules, these functional differences that affect T cells are undetermined. Especially, the differences in downstream signals and transcriptional activities under co-stimulatory molecules remained largely elusive and are even discrepancies. The discrepancy about differences in downstream co-stimulatory molecules mainly seems to be based on experimental conditions such as using CD28 and ICOS antibodies (Arimura et al., 2002; Wan et al., 2021) or co-culture with CHO cells which overexpress co-stimulatory molecules (Olsson et al., 1998; Slavik et al., 1999; Vasu et al., 2003). The agonistic antibodies for mimic co-stimulation are valuable to study the qualitative differences in CD28 and ICOS. However, activation states might not be equivalent to endogenous co-stimulation. In addition, although the experimental approaches using CHO cells might reflect physiological conditions of the interaction between T cells and antigen-presenting cells (APCs), the results are most likely to be interfered with unexpected interactions due to multiple regulation systems between co-stimulatory and co-inhibitory molecules. For example, although CD80 and CD86 bind to not only CD28 but also CTLA4 (Linsley et al., 1991b; Linsley et al., 1994), the interaction manners of CD80 and CD86 to CD28 and CTLA4 are distinct (Collins et al., 2002). Moreover, CD80, but not CD86 and ICOSLG, bind to PDL1 on APCs in cis-heterodimerization, escaping from T cell impairment through PD1 on T cell (Butte et al., 2007; Chaudhri et al., 2018; Sugiura et al., 2019; Zhao et al., 2019).

To exclude unexpected effects, we used CD80-Ig, CD86-Ig, and ICOSLG-Ig proteins for co-stimulations. In addition, we used synchronized culture cells EL4 as a homogeneous cell population. Here, we found remarkable differences in CD80, CD86, and ICOSLG. CD80-Ig + TCR stimulus promotes multiple gene expressions that are based on the long-term duration and massive activations of ERK and AKT pathways. These observations were positively correlated with binding activities of co-stimulatory molecules with T cells. Furthermore, the hyper-activated T cells by CD80-Ig + TCR stimulus were susceptible to disruption of effector activities by PDL1, due to promoted PD1 expressions through activated ERK and AKT. Our study provides insight into the underlying mechanism that co-stimulatory molecules play a pivotal role in deciding T cell fate through promoting co-inhibitory molecules such as PD1 besides T cell effector genes. These results shed light on molecular differences between co-stimulatory molecules.

## Results and discussion

### CD80 highly promotes expression of T cell effector genes in early phase, but induces expression of T cell exhaustion-related genes in late phase compared with CD86 and ICOSLG

To elucidate the functional differences in CD80, CD86, and ICOSLG, we conducted RNA-Seq for identifying expressed genes by these stimuli with anti-CD3 Ab (anti-CD3) as a mimic for TCR stimulation. Synchronized EL4 cells by cultured in 0.5 % FBS medium for 24 h were stimulated by co-stimulatory molecules with anti-CD3 (Fig. 1A). After stimulations, RNA samples were collected at 3, 8, and 24 h to evaluate gene expressions with time courses (Fig. 1A). Principal component analysis (PCA) revealed the qualitative differences in combined stimuli (Fig. 1B). A CD80-Ig + anti-CD3 stimulus was clearly discriminated from single anti-CD3, and CD86-Ig or ICOSLG-Ig + anti-CD3 stimuli. Next, we measured the transitions of differentially expression genes (DEGs) numbers induced by all stimuli. DEGs were defined by an absolute fold change (FC) > 2 and false discovery rate (FDR) < 0.05 compared with Ig treatment. CD80-Ig + anti-CD3 stimulus markedly amplified or suppressed a variety of genes (Fig. 1C and D). Venn diagram showed that the large part of DEGs by CD86-Ig or ICOSLG-Ig + anti-CD3 were included in DEGs induced by CD80-Ig + anti-CD3 stimulus, regardless of upregulated- or downregulated genes (Fig. 1E and F). Next, we conducted gene-set enrichment analysis (GSEA) to elucidate characteristics of altered expression genes in CD80-Ig + anti-CD3 stimulus compared with CD86-Ig or ICOSLG-Ig + anti-CD3 stimulus (Fig. 1G). As a result, although T cell activation-related gene sets were highly enriched at early time points, almost all of these gene sets were not significant at 24 h (q-value > 0.25). It suggested that effector activities decreased in CD80-Ig + anti-CD3 stimulus in a time-dependent manner. Additionally, by analysis using immunologic gene sets, T cell activation-related genes were significantly enriched at 3 h in CD80-Ig + anti-CD3 stimulus (Fig. 1H). By contrast, T cell exhaustion-related genes were highly enriched in CD80-Ig + anti-CD3 stimulus compared with in CD86-Ig or ICOSLG-Ig + anti-CD3 stimulus at 24 h (Fig. 1I).

**Fig. 1.**
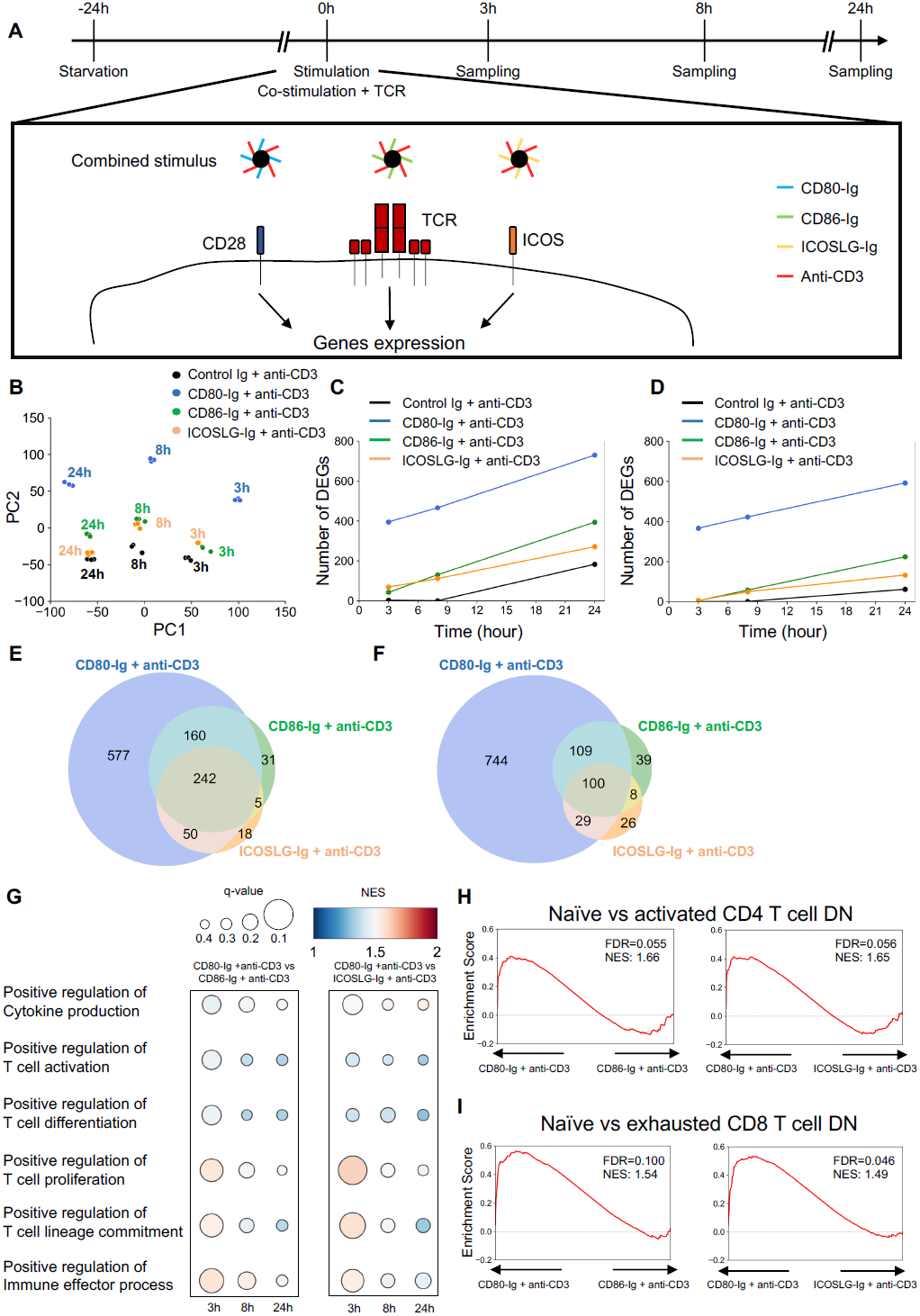
The difference of the regulated and enriched genes under co-stimulations with anti-CD3 (A) Experimental design for collecting RNA samples by each stimulus. (B) PCA analysis using mRNA expression data induced by co-stimulations + anti-CD3 stimuli. (C) Transitions of upregulated total DEG numbers compared with Control Ig. (D) Transitions of downregulated total DEG numbers compared with Control Ig. (E) Venn diagram of upregulated genes (FC >2 and FDR < 0.05). (F) Venn diagram of downregulated genes (FC <0.5 and FDR < 0.05). (G) GSEA based on ontology gene sets were conducted by comparing CD80-Ig + anti-CD3 with CD86-Ig or ICOSLG + anti-CD3 at each time point. The gene sets related to T cell effector activities were extracted and depicted based on q-value and normalized enrichment score (NES). The left square shows a comparison between CD80-Ig + anti-CD3 and CD86-Ig + anti-CD3. The right square shows a comparison between CD80-Ig + anti-CD3 and ICOSLG-Ig + anti-CD3. (H and I) GSEA based on immunologic signature gene sets were conducted by comparing CD80-Ig + anti-CD3 with CD86-Ig or ICOSLG + anti-CD3. At 3 h in H and 24 h in J. The outputted NES and q-value from the consequence of analysis were shown as enrichment plots.

### The expression levels of T cell effector genes and proteins are distinct from each combination of TCR with co-stimulatory molecules

Next, we examined expression intensities of representative genes that are categorized as cytokines and receptors, TNF superfamily, transcription factors, and co-inhibitory molecules (Fig. 2A). The genes related to effector functions and T cell exhaustion were prominently enhanced by CD80-Ig + anti-CD3 stimulus compared with other combined stimuli. These results demonstrated that not only the number of DEG but also the amount of change was remarkably fluctuated in CD80-Ig stimulus compared with CD86-Ig or ICOSLG-Ig. Next, we conducted temporal expression analysis of transcription factors, cytokines, and co-inhibitory molecules. As the representative genes, Batf, Irf4, Arnt2, Pdcd1, Vsir, and Tnf were shown to change over time from baseline (Fig. 2B-G). Transcription factors Batf and Irf4 mRNA were upregulated at early time points, probably due to involving regulation of T cell activation as key transcription factors (Fig. 2B and C). Arnt2 mRNA was enhanced in a time-dependent manner by all combined stimuli (Fig. 2D). Pdcd1 and Vsir mRNA was gradually upregulated by all stimuli (Fig. 2E and F). Tnf mRNA was shown to peak at 3 h and was rapidly attenuated by CD80-Ig + anti-CD3 stimulus (Fig. 2G). In all genes, CD80-Ig + anti-CD3 stimulus highly induces gene expression compared with other combined stimulations at all time points. To evaluate whether the temporal patterns of these mRNA were consistent with protein levels, we examined protein expressions by immunoblotting (Fig. 2H). As corresponding with expressing transitions of mRNA, transcriptional factor IRF4 and ARNT2 was enhanced from at 3 h and 8 h, respectively. However, temporal patterns of BATF protein expressions were not consistent with that of Batf mRNA expressions. Intriguingly, CD80-Ig + anti-CD3 stimulus upregulated PD1 from 24 h, but dual ICOSLG-Ig + anti-CD3 stimuli enhance PD1 protein levels at 48 h. In CD80-Ig or CD86-Ig + anti-CD3 stimulus, VISTA levels were enhanced at 24 h (Fig. 2H). We evaluated the expression levels of cytokines in the supernatants by a cytokine array (Fig. 2I and J). CD80-Ig + anti-CD3 stimulus promoted higher secretions of cytokines that are involved in T cell activations (Fig. 2I). Each spot was quantified for visualization by Heatmap. CD80-Ig + anti-CD3 stimulus remarkably promoted secretions of almost all cytokines compared with Control Ig, CD86-Ig, or ICOSLG-Ig + anti-CD3 (Fig. 2J).

**Fig. 2.**
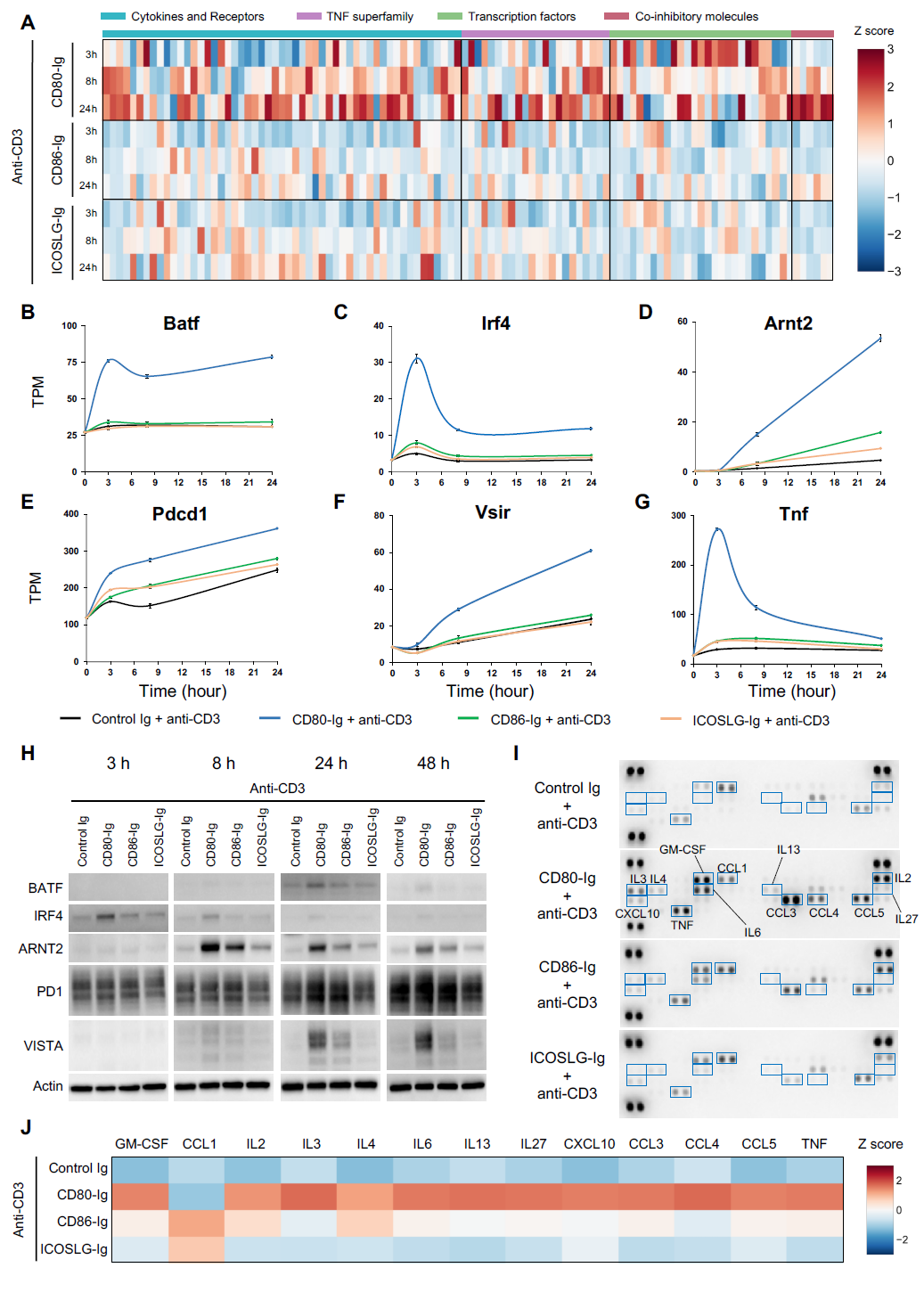
The intensity of gene and protein expressions that are related to effector activities and T cell impairment are distinct in the combination of co-stimulations with anti-CD3 (A) Heatmap of Z scores shows genes that are categorized in cytokines and receptors, TNF superfamily, transcription factors, and co-inhibitory molecules. (B-G) The chronological changes of mRNA expression levels from baseline. Batf in B, Irf4 in C, Arnt2 in D, Pdcd1 in E, Vsir in F, and Tnf in G. (H) Altered expressions of protein with time transitions. CD80-Ig + anti-CD3 stimulus enhanced BATF, IRF4, ARNT2, PD1, and VISTA compared with single anti-CD3, CD86-Ig or ICOSLG-Ig + anti-CD3. (I and J) The expression levels of cytokines induced by single anti-CD3 or individual co-stimulation + anti-CD3 in supernatants were measured by the cytokine array. Acquired images were shown in I. Quantified counts of expression levels were normalized and visualized as a heatmap of Z scores in J.

### Co-stimulatory molecules regulate activation levels of ERK and AKT, which are critical for the expressions of T cell effector proteins

To elucidate the underlying regulation mechanisms of DEG numbers and their expression levels by each co-stimulatory molecule, we monitored phosphorylation status of ERK and AKT which are primary mediators of TCR and co-stimulations. All combined stimuli promoted ERK phosphorylation from 5 min (Fig. 3A). Phosphorylated AKT was observed by all combined stimuli excluding Control Ig + anti-CD3 stimulus (Fig. 3A). To compare the activation status with stimulations in time courses, the observed bands of phosphorylated ERK and AKT were quantified and normalized by expression levels of Actin. Compared with CD86-Ig or ICOSLG-Ig + anti-CD3 stimuli, CD80-Ig + anti-CD3 stimulus showed remarkable activation of ERK and AKT with high magnitude and long-term (Fig. 3B and C). Interestingly, the amplified and sustained ERK and AKT under co-stimulatory molecules + anti-CD3 stimuli were correlated with DEGs numbers (Fig. 1E and F), suggesting that the variety of gene expression induced by CD80-Ig + anti-CD3 was based on a consequence of strong induction of downstream pathways.

**Fig. 3.**
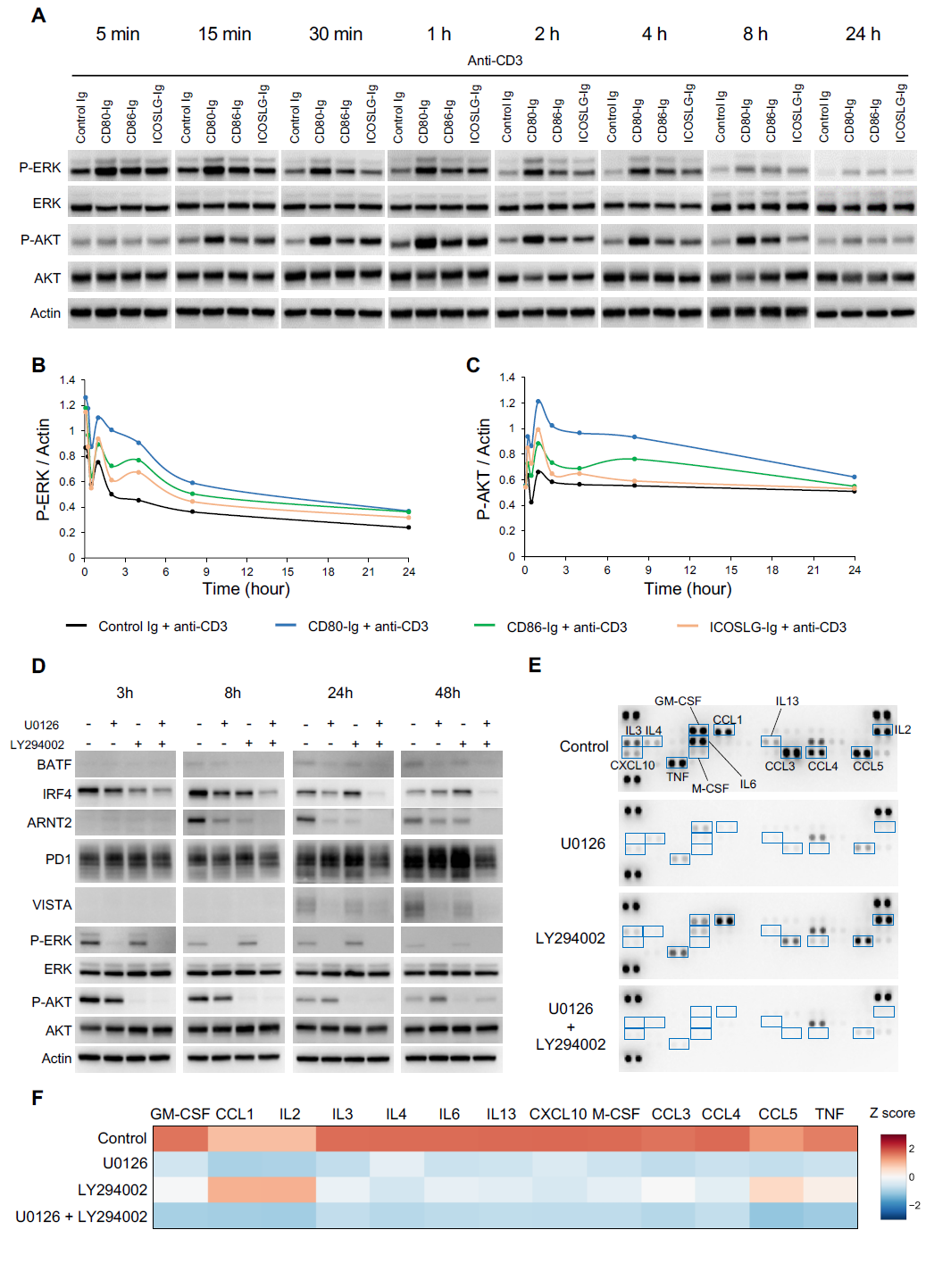
The magnitudes and durations of activated ERK and AKT under co-stimulations + anti-CD3 are key mediators for expressions of T cell activations- and exhaustions-regulated genes (A) Immunoblotting for the chronic changes of phospho-ERK and phospho-AKT in EL4 cells stimulated by anti-CD3 with or without a co-stimulation. (B and C) Transitions of activated status were shown. Captured images of the phospho-ERK or phospho-AKT bands were quantified. Those counts were normalized by quantified counts of Actin bands. The changes of phospho-ERK in B and that of phospho-AKT in C. (D) The effects on protein expressions by single or simultaneous inhibitions of ERK and AKT. 10 μM of U0126 and LY294002 were pretreated before 1 h from stimulation. Cell lysates were collected at indicated time points after stimulation. (E and F) The effects of single or simultaneous inhibitions of ERK and AKT on expression levels of cytokines induced by CD80-Ig + anti-CD3 in supernatants were measured by the cytokine array. 10 μM of U0126 and LY294002 were pretreated before 1 h from stimulation. After 24 h of stimulation, the supernatant was collected. Acquired images were shown in E. Quantified counts of expression levels were normalized and visualized as a heatmap in F.

Next, we examined the effects to protein expression in condition with limited activations of ERK and AKT under CD80-Ig + anti-CD3 stimulus. Synchronized EL4 cells were pre-treated with a MEK inhibitor U0126 or a PI3K inhibitor LY294002 for 1 h and they were stimulated by CD80-Ig + anti CD3 stimulus. Cell lysates were collected after 3, 8, 24, 48 h from stimulations and were conducted immunoblotting. We confirmed suppressions of phospho-ERK and phospho-AKT by U0126 and LY294002 treatment (Fig. 3D). Although BATF, IRF4 and ARNT2 expressions were partially abrogated by a single inhibition of ERK or AKT, they were completely suppressed by dual inhibition (Fig. 3D). PD1 expressions were not inhibited by a single inhibition, but dual inhibitions suppressed upregulation of PD1 by CD80-Ig + anti-CD3 stimulus at 24 and 48 h. VISTA expressions were adequately suppressed by a single inhibition of ERK activities (Fig. 3D), but abrogated AKT activity showed partial effects on VISTA expression at 48 h. These data demonstrate that ERK and AKT signaling regulate gene expressions with independently, cooperatively, or redundantly. Moreover, co-inhibitory molecules such as PD1 and VISTA were regulated downstream of ERK and AKT pathways activated by robust stimuli. Next, we evaluated the effects on cytokine expressions by protein array under restricted conditions of the ERK and AKT signaling (Fig. 3E) and expression levels were quantified (Fig. 3F). Suppression of ERK by U0126 downregulated secretion of almost all cytokines (Fig. 3E and F). By contrast, inhibitions of AKT showed limited suppression of the cytokine compared with U0126 treatment (Fig. 3E and F). The simultaneous inhibition of ERK and AKT completely suppressed cytokine expressions despite CD80-Ig + anti-CD3 stimulus (Fig. 3E and F). These results suggest that the magnitude and duration of ERK and AKT activations that are enhanced by combined stimuli is essential for effective activations of T cells and following to T cell exhaustions. In addition, the ERK and AKT pathways synergistically promotes protein expressions under the co-stimulatory molecules and TCR.

### The binding activities to T cells are distinct in co-stimulatory molecules

To elucidate the underlying mechanism of deciding signal intensity and duration downstream of each co-stimulatory molecule, we measured binding activities between co-stimulatory molecules and EL4 cells by cell isolation assays. Magnetic beads conjugated Ig proteins with and without anti-CD3 were reacted with intact EL4 cells, and then the number of isolated cells was quantitatively measured (Fig.S1A). CD80-Ig intensely engaged with EL4 cells compared with CD86-Ig and ICOSLG-Ig (Fig. 4A). The order of binding activities by co-stimulation + anti-CD3 was consistent with the case of cell separation by single co-stimulatory molecules (Fig. 4B). ICOSLG-Ig showed weak binding activities to EL4 cells regardless of with and without anti-CD3. Next, we established CD28 or ICOS knockout cells by CRISPR-Cas9 editing and evaluated binding activities between co-stimulatory molecules and knockout cells. We observed indels closed to the target regions of gRNA in both knockout cell lines (Fig. S1B and S1C) and confirmed complete ablation (Fig. 4C and D). In CD28 knockout cells, binding activities of CD80-Ig and CD86-Ig were completely disrupted compared with parental cells (Fig. 4E and F). In ICOS knockout cells, binding activities of ICOSLG-Ig were also entirely disrupted compared with parental cells (Fig. 4G). These results indicate that the targets of CD80 and CD86, or ICOSLG are only CD28 or ICOS on EL4 cell line. And then, we performed immunoprecipitation assay to verify interaction CD28 on EL4 cells with CD80-Ig and CD86-Ig or ICOS on them with ICOSLG-Ig (Fig. 4H). CD80-Ig strongly interacted with CD28 in EL4 lysates. However, CD86-Ig showed weaker affinity with CD28 than CD80-Ig. ICOSLG-Ig showed efficient interaction with ICOS, but not with CD28. Regarding CD80 and CD86, these results seemed to reflect differences in cell isolation efficiencies (Fig. 4A and B). From these results, we revealed the binding activities to EL4 cells are positively correlated with the magnitude and duration of activated ERK and AKT signals and total DEG numbers, suggesting that the binding strength of co-stimulatory molecules to T cells is a critical factor for deciding activities of their downstream signaling and diversity of gene expressions.

**Fig. 4.**
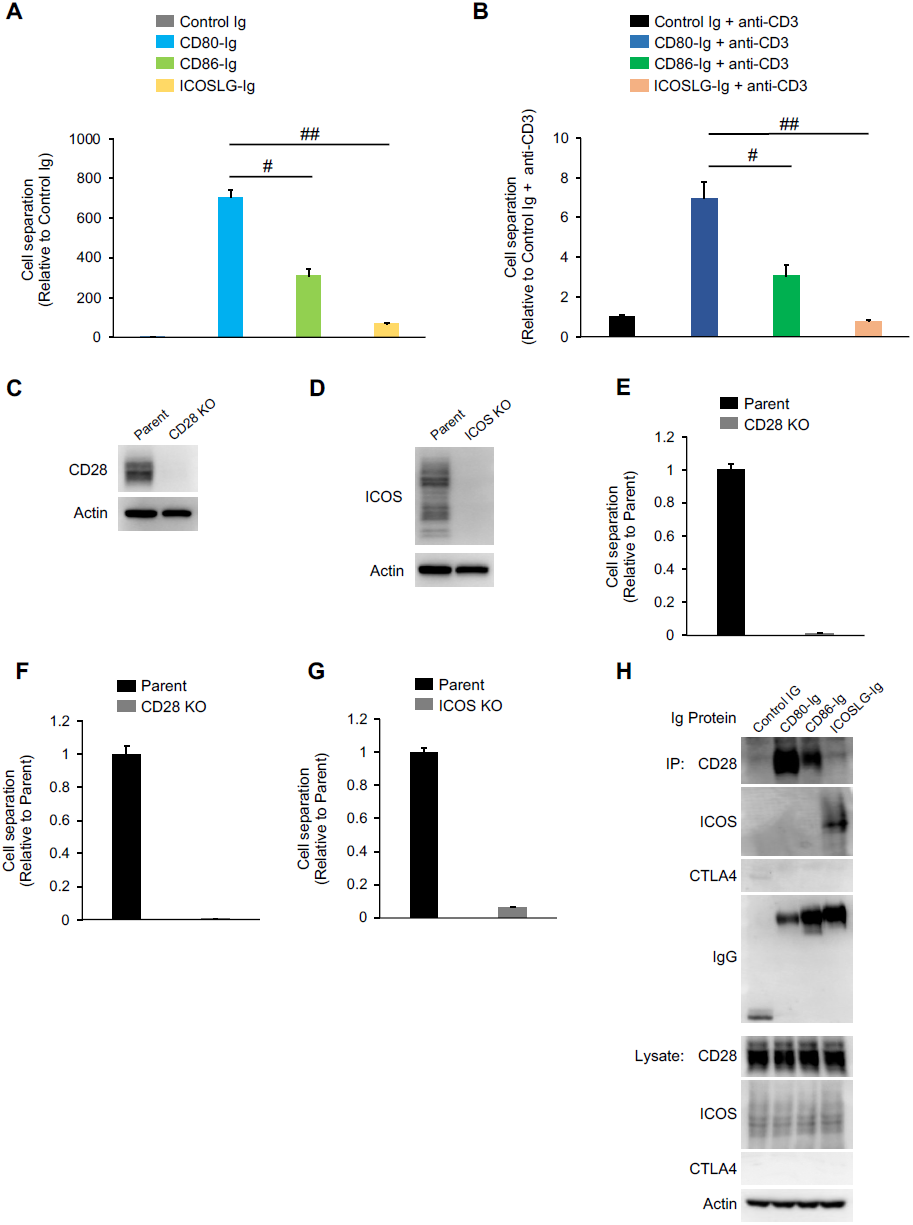
Binding activities of co-stimulatory molecules to T cells (A) The binding intensities of co-stimulatory molecules with EL4 cells were measured. Shown is the ratio of the binding intensities by Control Ig. Data are means ± SEM from two independent experiments performed in quadruplicate. #, p < 0.05; ##, p < 0.05. (B) The binding intensities of co-stimulatory molecules + anti-CD3 with EL4 cells were measured. Shown is the ratio of the binding intensities by Control Ig + anti-CD3. Data are means ± SEM from two independent experiments performed in quadruplicate. #, p < 0.05; ##, p < 0.05. (C and D) CD28 or ICOS expressions were impaired by CRISPR-Cas9-dependent genome editing. In C, ablation of CD28 proteins in CD28 KO cells. In D, ablation of ICOS protein in ICOS KO cells. (E and F) The binding intensities of CD80 (E) or CD86 (F) with EL4 cells were measured. Shown is the ratio of the binding intensities to CD80-Ig (E) or CD86-Ig (F) by Parent cells. Data are means ± SEM from two independent experiments performed in quadruplicate. (G) The binding intensities of ICOSLG with EL4 cells were measured. Shown is the ratio of the binding intensities to ICOSLG-Ig by Parent cells. Data are means ± SEM from two independent experiments performed in quadruplicate. (H) Co-immunoprecipitation of CD80-Ig, CD86-Ig, or ICOSLG-Ig with CD28, ICOS, or CTLA4 in EL4 lysates.

### Hyper activated T cells are vulnerable to PDL1 and are susceptible to suppressions of T cell effector genes and proteins

Based on the enrichment of T cell exhaustion-related genes in CD80-Ig + anti-CD3 stimulus (Fig. 1I), we examined the inhibitory effect of PD1:PDL1 engagements on gene expression after combined stimuli by RNA-Seq. First, we testified to the differences in PD1:PDL1 binding activitie. After 24 h from the combined stimulations, we collected EL4 cell lysates and performed co-immunoprecipitation using Control Ig and PDL1-Ig. Due to slight PD1 amplifications by the massive stimulation, PDL1-Ig effectively immunoprecipitated PD1 in the lysates from CD80-Ig + anti-CD3 stimulus compared with other combined stimuli (Fig. 5A). For RNA-Seq analysis, synchronized EL4 cells cultured in 0.5 % FBS medium for 24 h were activated by combined stimuli. After 24 h from the combined stimulations, Control Ig or PDL1-Ig was added to stimulated EL4 cells, and total RNA samples were collected after further 24 h (Fig. 5B). To classify all samples based on expression genes, we conducted PCA (Fig. 5C). Although the cluster of stimulated cells by CD80-Ig + anti-CD3 → PDL1-Ig was distinct from that of the CD80-Ig + anti-CD3 → Control Ig, the differences in CD86-Ig or ICOSLG-Ig + anti-CD3 stimulus between Control Ig and PDL1-Ig were limited (Fig. 5C). Here, we defined DEGs as FC < 0.5 and FDR < 0.05 in comparison between Control Ig and PDL1-Ig. Venn diagram of DEG number showed that suppressed DEGs by PDL1-Ig were numerous in CD80-Ig + anti-CD3 stimulated cells compared with CD86-Ig or ICOSLG-Ig + anti-CD3 stimulated cells (Fig. 5D). To evaluate vulnerabilities to PDL1 on induced genes by each combined stimuli, we examined the fluctuations of DEGs categorized with cytokines and receptors, transcription factors, and co-inhibitory molecules. Despite excessive expressions, the induced mRNAs by CD80-Ig + anti-CD3 were highly sensitive to suppressions of PDL1 compared with other combined stimuli (Fig. 5E). We evaluated expression levels of cytokines after Control Ig or PDL1-Ig by cytokine array. Consistent with the results of RNA-Seq, the cytokine induced by CD80-Ig + anti-CD3 stimulus was highly sensitive to suppressions by PDL1-Ig compared with other combined stimulations (Fig. 5F and Fig. S2). These data demonstrated that intensely activated T cells are relatively sensitive to co-inhibitory molecules. Finally, we quantified obtained results as four categories in binding activities to T cells, ERK and AKT activities, induced gene numbers, and vulnerable gene numbers to PDL1. Quantified data were relatively compared between combined stimuli in each category. As a result, the binding activities of co-stimulatory molecules to T cells positively correlated with the activation status of ERK and AKT, altered DEG numbers, and suppression effector gene numbers by PDL1 (Fig. 5G). These results showed that co-stimulatory molecules govern not only the T cell effector activities but also vulnerabilities to PDL1 through regulations of ERK and AKT activations, demonstrating that the mechanism plays a key role in deciding T cell fate (Fig. 5H).

**Fig.5.**
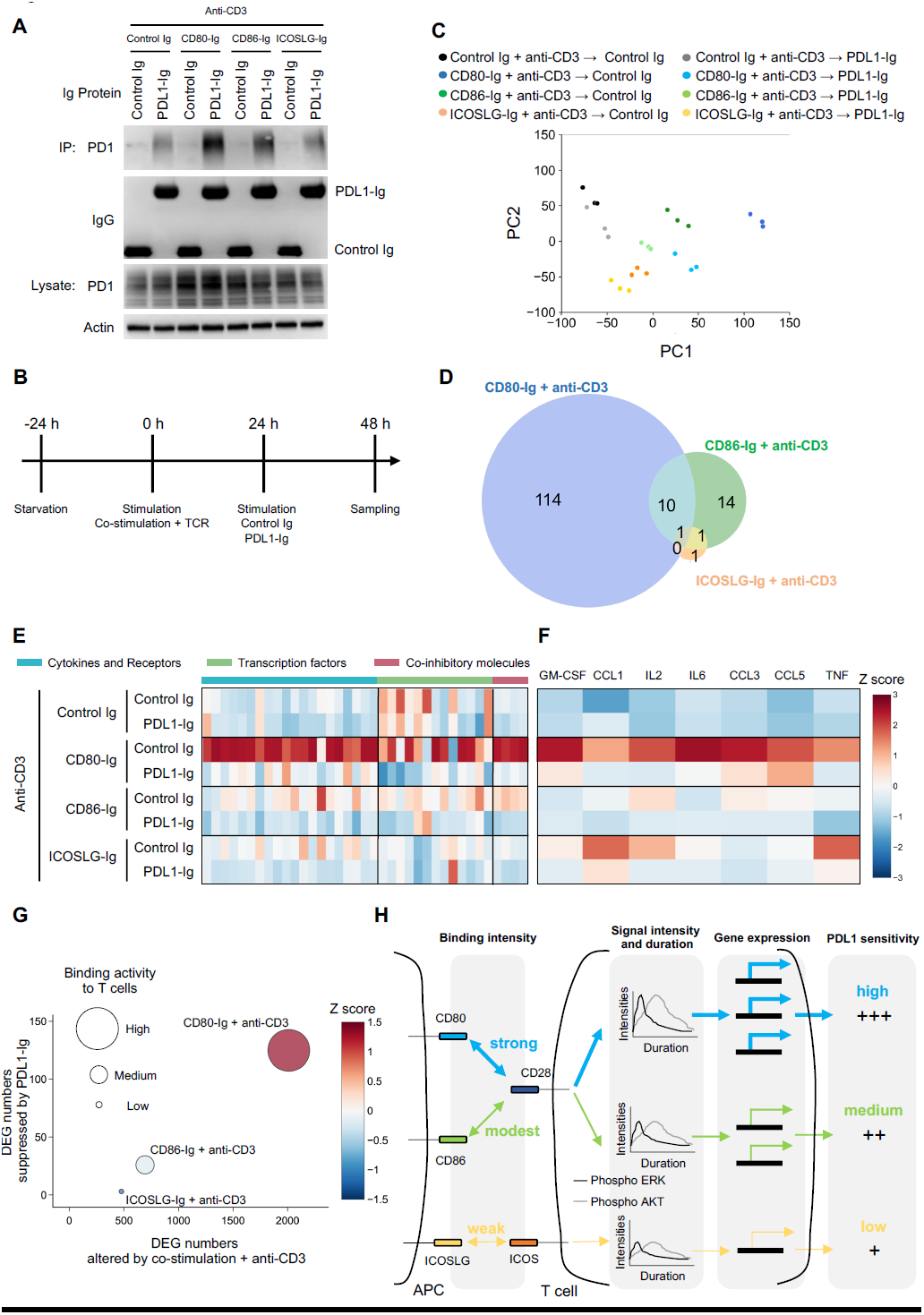
T cell vulnerabilities to PDL1 are distinct depending on the pretreated type of co-stimulation. (A) The interactions of PD1 that were induced by individual stimuli with PDL1-Ig were evaluated by a co-immunoprecipitation assay. (B) Experimental design for collecting RNA samples by each stimulus. (C) PCA analysis was conducted using mRNA expression data from combined stimulations →Control Ig or PDL1-Ig stimulus. (D) Venn diagram of DEG (FC < 0.5 and FDR < 0.05) compared between control Ig and PDL1-Ig after the combined stimulus. (E) Heatmap of Z scores shows the fluctuations of DEGs that were related to cytokines, transcription factors, and co-inhibitory molecules. (F) The effects of PDL1-Ig to expression levels of cytokines induced by single anti-CD3 or combined stimulus in supernatants were measured by the cytokine array. Acquired images that are shown in Fig. S2 were quantified, normalized, calculated Z score, and visualized as a heatmap. (G) Bubble plot shows the comparison of quantified counts in each index generated by combined stimulus. The total under the curve of Phospho-ERK and phospho-AKT in Fig. 3B and C were calculated and compared between combined stimuli for showing as a Z score. Bubble size shows binding activities to T cells that is related to Fig. 4B. The horizontal axis shows observed DEG number by combined stimulus that is related to Fig. 1H and 1I. The vertical axis shows suppressed DEG number by PDL1 that is related to Fig. 5D. (H) The schematic image of the correlation between binding activities and T cell vulnerability to PDL1. Binding activities to T cells, the duration and intensity of ERK and AKT, induced genes, and sensitivities to PDL1 are decided by the type of co-stimulation, and these indexes are positively correlated.

The differences between CD80, CD86, and ICOSLG were uncertain and inconsistent for several decades. Here, we investigated the functional differences of CD80, CD86, and ICOSLG in T cell invigorations and exhaustions using Ig proteins and a uniform cell population for avoiding unexpected interference. We revealed that quantitative differences in avidity with T cells caused by co-stimulatory molecules positively associated with the magnitude and duration of ERK and AKT phosphorylation and following the variation of effector genes expression (Fig. 5H). Additionally, we elucidated that hyper-activated ERK and AKT induce the expression of co-inhibitory molecules, thereby preferentially suppressing T cell activities. Indeed, in response to CD80 + TCR stimulus, T cells intensively expressed effector genes at the early time point and induced co-inhibitory molecules including PD1 at the late time point (Fig. 2H-J). However, T cells stimulated by CD86 or ICOSLG + TCR stimuli are relatively resistant to PDL1 stimuli instead of the limited effector activities. The bias seems reasonable to disrupt and exclude aberrantly activated T cells for avoiding excessive inflammations in physiological conditions. These functional differences of co-stimulatory molecules in the regulations of T cell effector activities and impairments might be one of the biological significance to exist multiple co-stimulatory molecules that are involved in T cell activation.

The duration and magnitude of ERK activities regulate cell fate decisions in PC12 cells (Ebisuya et al., 2005; Marshall, 1995). Treatment of PC12 cells with NGF causes differentiation into sympathetic-like neurons by the sustained ERK activations, whereas treatment with EGF promotes proliferation by transient ERK activations. Further research elucidated that the association between phospho-ERK and phospho-AKT response separates differentiation from proliferation in PC12 cells (Chen et al., 2012). However, it remained unclear whether both serine/threonine kinases control the T cell fates. In the present study, we elucidated that the duration and magnitude of active ERK and AKT generated by co-stimulatory molecules affect different cellular responses such as effector activities and vulnerabilities to PDL1 ligations (Fig. 5H). Moreover, from the limited inductions of effector and co-inhibitory molecules by simultaneous inhibitions of ERK and AKT (Fig. 3D-F), both serine/threonine kinases cooperatively regulate T cell invigoration and impairment. These observations are strong evidence that ERK and AKT are decision-makers for T cell fates. It is assumed that transcriptional factors NFAT, IRF4, and BATF are especially key molecules downstream of ERK and AKT for T cell fate decisions, due to promote expressions of cytokines and PD1 (Dienz et al., 2007; Man et al., 2017; Martinez et al., 2015). Indeed, expression levels of these transcriptional factors differ depending on co-stimulation (Fig. 2B and C). Therefore, consistent with PC12 cells in response to EGF or NGF, the variations of ERK and AKT activities in T cells upon stimulations by co-stimulatory molecules cause distinct cellular responses. These mechanisms might play a critical role in generating heterogeneous T cell populations for the effective eradication of pathogens and cancer cells.

Multiple co-stimulatory and co-inhibitory molecules other than CD80, CD86, and ICOSLG orchestrate the effector activities of T cells (Attanasio and Wherry, 2016; Chen and Flies, 2013; Wykes and Lewin, 2018; Zhang and Vignali, 2016). However, most of them about the functional redundancies and differences remain to be elucidated. The investigation of whole underlying mechanisms in T cell invigoration and impairment is a key step for understanding T cell biology. Additionally, the challenges shed light on developing and improving CAR-T therapies and immunotherapies. We can observe specific signals and gene expression by stimulations using Ig protein. The method will be broadly applied for a comprehensive understanding of the activated signaling pathways and gene expressions in T cell regulations by co-stimulatory and co-inhibitory molecules.

## Materials & methods

### Cell culture

EL4 parent, EL4 CD28 KO, EL4 ICOS KO, or Ig protein expressing MC38 cells were maintained RPMI1640 supplemented with 10% Fetal Bovine Serum (FBS). For synchronization, EL4 cells were cultured in RPMI1640 supplemented with 0.5% FBS for 24 h. All cell cultures were maintained in an incubator at 37 °C with 5% CO_2_.

### Purification of Ig Proteins

CD80-Ig, CD86-Ig, ICOSLG-Ig, and PDL1-Ig were designed as chimeric proteins that are composed of the ectodomain of CD80, CD86, ICOSLG, and PDL1 with IgG4 Fc domain. MC38 was transferred these plasmids coding each Ig proteins for stable supplying Ig proteins. For collections of Ig proteins, engineered MC38 expressing endogenously Control Ig, CD80-Ig, CD86-Ig, ICOSLG-Ig, or PDL1-Ig were cultured in RPMI1640 supplemented with 10% Ultra Low IgG FBS (GIBCO). Culture supernatants were collected and extracted by Sepharose beads (GE Healthcare) for co-immunoprecipitation or magnetic beads (Thermo Fisher) for stimulation and cell isolation assays. An equal volume of Ig proteins was used for each experiment.

### Knockouts of CD28 or ICOS and indel identifications

CRISPR Cas9 systems were applied for the establishment of CD28 or ICOS knockout cells. gRNA (CD28: TCGGCATTCGAGCGAAACTG, ICOS: AGGTTCCTTTCTTGAAAAGG) with Cas9 protein (Thermo Fisher) was incubated for 5 min, and the mixtures were introduced into EL4 parent cells by electroporation. For indel identification, extracted genomic DNAs from EL4 parental, CD28 KO, and ICOS KO cells were amplified regions that are close to target sequences for gRNA. Primers were designed as follow: CD28 Forward: CTAACATCATAGGAATCTCA, CD28 Reverse: TGTCAGACTATGTTTCATTG or ICOS Forward: TGTCTATCACATGAAAAGCC, ICOS Reverse: GAAACTTTACGAATCTAGTG. To allow mismatched DNA to form, PCR products were denatured and re-annealed. The re-annealed PCR products were reacted with T7 endonuclease I (NIPPONGENE) for 30 min at 37 °C and the reactants were conducted gel analysis by agarose electrophoresis.

### Co-stimulations with anti-CD3 and pretreatments with inhibitors

Synchronized EL4 cells were stimulated by Ig Protein with anti-CD3 (final 2μg/ml, Biolegend) conjugated magnetic beads. RNA samples and protein samples from EL4 cells and culture supernatants were harvested for RNA-Seq, immunoblotting, and cytokine arrays at indicated time points. For inhibition of ERK or AKT signal, synchronized EL4 cells were pretreated with 10μM U0126 or LY294002 for 1 h before stimulation, and then cells were stimulated by prepared magnetic beads.

### RNA-Seq analysis

For samples preparation, stimulated EL4 cells were harvested at indicated time points and total RNA were extracted by RNeasy Mini kits (Qiagen) according to the manufacturer’s instructions. mRNA sequencing libraries were prepared with 500 ng of total RNA of each sample using the NEBNext Ultra II Directional RNA Library Prep Kit for Illumina (NEB) with NEBNext Poly(A) mRNA Magnetic Isolation Module (NEB), according to manufacturer’s protocol. Quality controls of the final library were performed using 2100 Bioanalyzer Instrument (Agilent). Paired-end sequencing was performed on Illumina NovaSeq 6000 (2 × 150 bp reads). The samples were multiplexed and sequenced on one High Output Kit v2.5 to reduce a batch effect.

For RNA-Seq analysis, BCL raw data was converted to FASTQ data and demultiplexed by bcl2fastq Conversion Software (Illumina). Raw reads for individual samples were assessed with the FastQC program to check sequence quality and processed with Trimmomatic to remove adaptor sequences and bases with low quality scores. Reads were aligned to the GRCm38/mm10 mouse reference genome using STAR. The number of reads mapped to each gene was counted by htseq-count version. Gene expression levels were quantified as transcripts per million using RSEM.

### GSEA

GSEA was conducted using whole transcriptome data. The ontology gene sets and immunologic signature from MSigDB was applied for Gene sets database. The RNA-Seq data of CD80-Ig + anti-CD3 were compared with CD86-Ig or ICOSLG-Ig + anti-CD3. Enrichment plots were depicted based on “Rank in gene list” and “Running enrichment score”.

### Antibodies

The following antibodies were used in this study: Actin (Cell Signaling, 4970), CD28 (Cell Signaling, 38774), ICOS (Cell Signaling, 67223), CTLA4 (Abcam, ab237712), IgG (Millipore, MAB1312), BATF (Cell Signaling, 8638), IRF4 (Cell Signaling, 62384), ARNT2 (MyBioSource, MBS2525431), PD1 (Cell Signaling, 84651), VISTA (Cell Signaling, 54979), P-ERK (Cell Signaling, 4370), ERK (Cell Signaling, 4695), P-AKT (Cell Signaling, 4060), AKT (Cell Signaling, 4691). Anti-rabbit IgG, HRP-linked Antibody (Cell Signaling, 7074) and Anti-mouse IgG, HRP-linked Antibody (Cell Signaling, 7076) were used as secondary antibodies.

### Immunoblotting

Cells were lysed by a lysis buffer (Lysis buffer 20 mM Tris-HCl at pH 7.5, 150 mM NaCl, 1 mM EDTA, 1 mM EGTA, and 1% Triton with proteinase inhibitors and phosphatase inhibitors). Protein concentration was determined with Bradford Protein Assays (Bio-Rad) and boiled in SDS sample buffer. 10 μg proteins were conducted SDS-PAGE electrophoresis, and the resolved proteins were then transferred to PVDF membrane (Millipore). The membrane was blocked using 5% skimmed milk and then immunoblotted with primary antibodies at 4°C overnight. After washing with 1xTBS-T, secondary antibodies were added and incubated at room temperature for 1 hour. Blot bands were detected with Immobilon (Millipore).

### Co-immunoprecipitation assay

Collected supernatant of engineered MC38 cells were purified Ig proteins by Sepharose beads. Prepared cell lysates were added to Sepharose beads conjugated with Ig proteins, and the reactions were conducted at 4 °C overnight. After washing the Protein A Sepharose beads by the same condition with above lysis buffer, and boiled in SDS sample buffer. The following procedure was same of the Immunoblotting.

### Cytokine array

Secreted cytokines from stimulated EL4 cells into cultured supernatants were detected using a Mouse Cytokine array kit (R&D) according to the instructions. For comparison of cytokine expressions between stimulations in the individual figure, images were obtained at the same time. Obtained signals were identified by transparency overlay, and spots were quantified using ImageJ software.

### Cell isolation assay

Purified Ig proteins with magnetic beads were added to tubes. Cultured EL4 parent, CD28 KO, or ICOS KO cells were gently washed by PBS, and equivalent numbers of EL4 parent, CD28 KO, or ICOS KO cells were added into tubes that contained magnetic beads. The mixtures were mildly suspended by pipetting and incubated for 20 min at 4 °C. For washing magnetic beads, tubes were set sample racks and washed three times by wash buffer (PBS including 2 mM EDTA and 0.5 % BSA). Captured cells were quantified by CellTiter-glo (Promega) and measured by Envision (PerkinElmer).

### Measurements of band intensities

The obtained signals by immunoblotting or cytokine arrays were performed using ImageJ software. For Fig. 3B and C, quantified the intensities of each signal were normalized by that of Actin band at the same lane. For Fig. 2J, Fig. 3F, and Fig. 5F, signals from cytokine arrays were quantified. Heatmaps showed Z scores that were based on the expression levels of cytokines comparison between co-stimulations or treatments with inhibitors.

### Statistical analysis

All data are presented as means ± standard error of the mean (SEM). The differences between the means of groups were analyzed by unpaired Student’s t-test or one-way analysis of variance (ANOVA) followed by Dunnett’s multiple comparison test. P values less than 0.05 were considered statistically significant.

## Acknowledgements

This work was supported by True Cell Simulations.

## Author contributions

Ken. I. designed and performed the experiments. Ken. I. and Kaz. I. wrote the manuscript. Ken. I. supervised the project.

---

**Fig S1.**
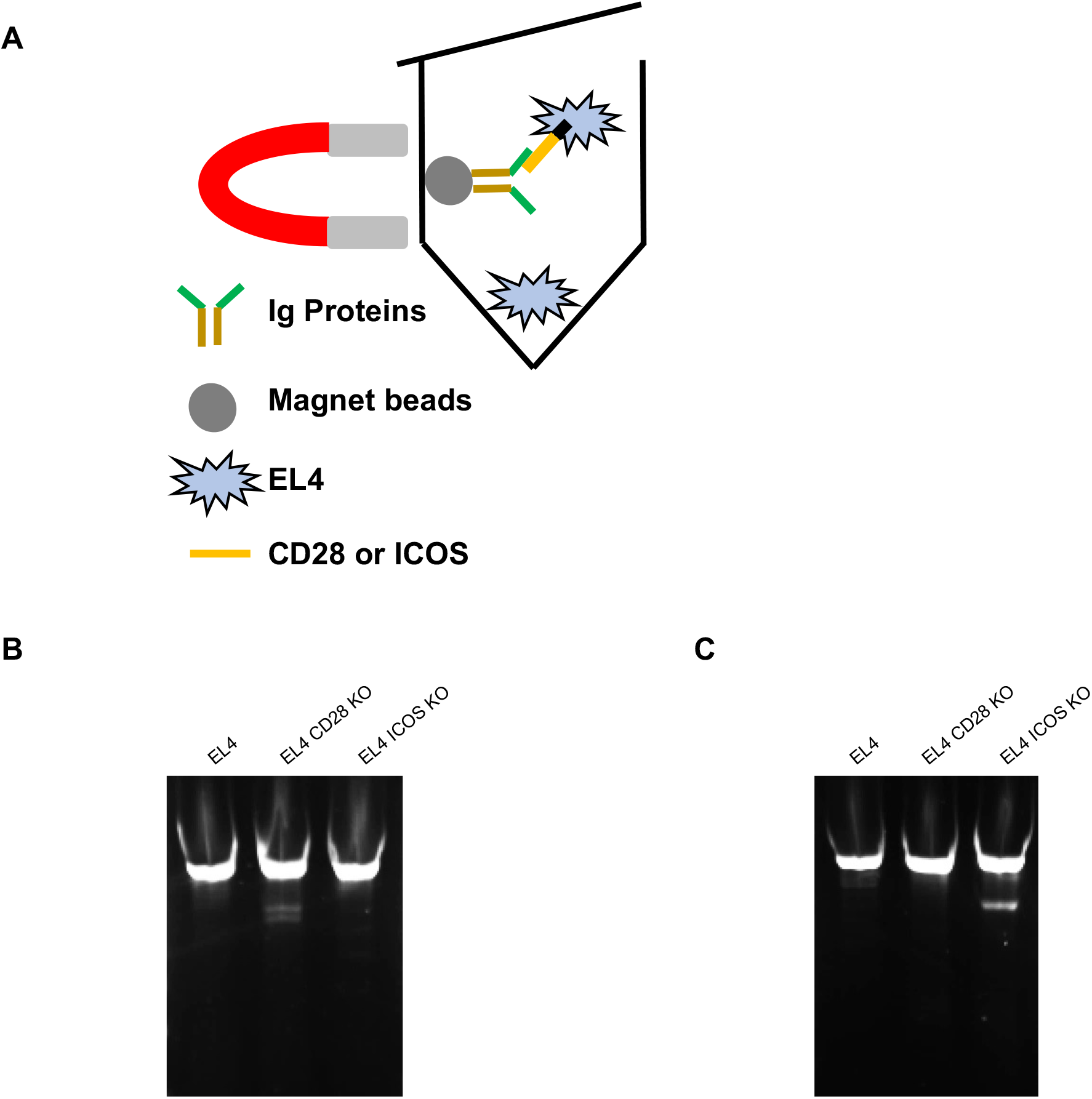
Cell separation assay and confirmations of CD28 or ICOS KO EL4 cells. (**A**) A schematic image of cell isolation assay. (**B and C**) Agarose gel electrophoresis were conducted using PCR products treated with T7 endonuclease I. In B, the regions of close to designed gRNA on CD28 were amplified. In C, the regions of close to designed gRNA on ICOS were amplified.

**Fig S2.**
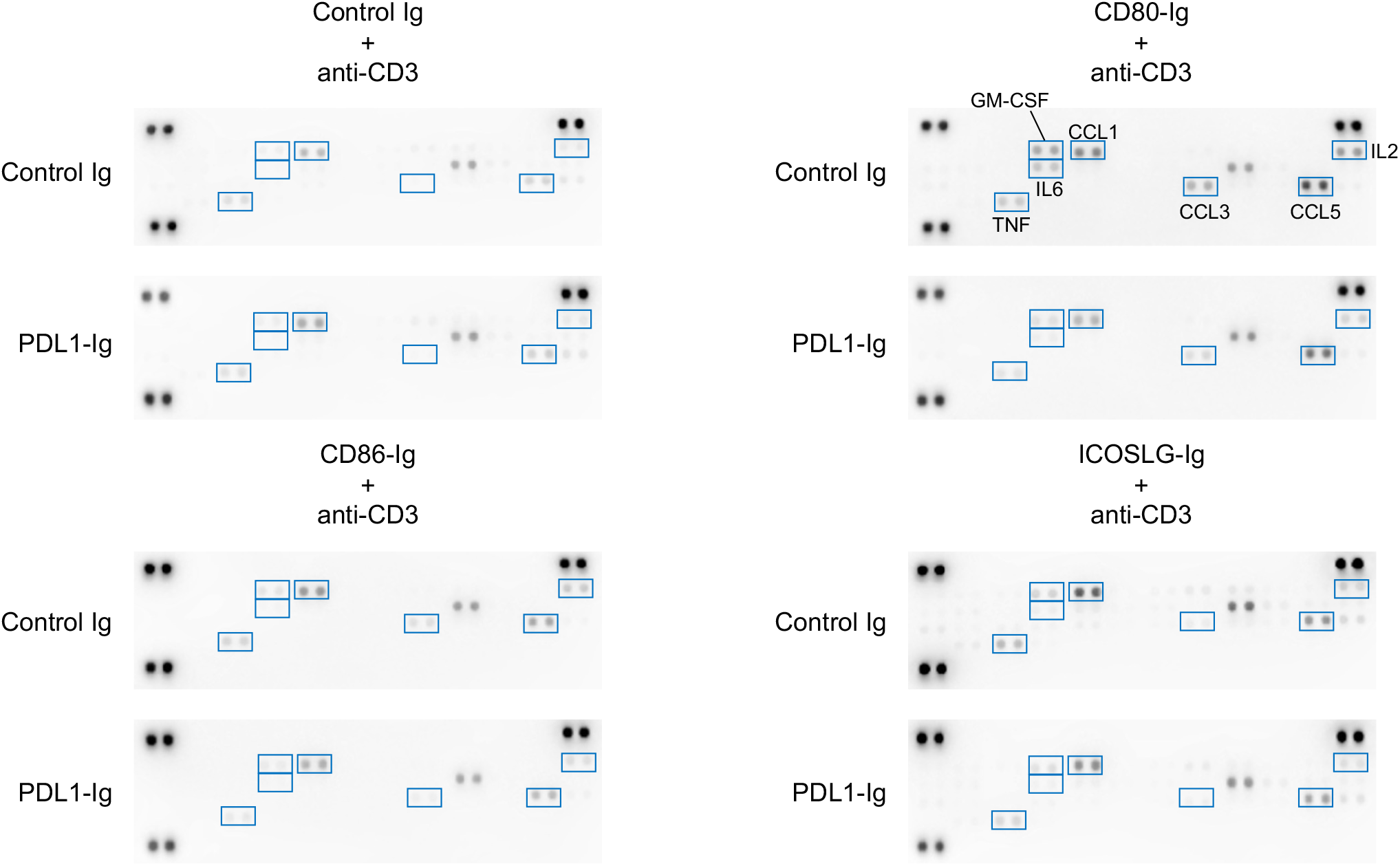
The relationships between co-stimulations and vulnerabilities to PDL1. The effects of PDL1-Ig to expression levels of cytokines induced by single anti-CD3 or combined stimulus in supernatants were measured by the cytokine array. Acquired images were shown. Quantified counts of expression levels were normalized, and visualized as a heatmap shown in Figure. 5F.

